# mmContext: an open framework for multimodal contrastive learning of omics and text data

**DOI:** 10.64898/2025.12.08.692934

**Authors:** Jonatan Menger, Sonia Maria Krissmer, Clemens Kreutz, Harald Binder, Maren Hackenberg

## Abstract

**Summary:** Multimodal approaches are increasingly leveraged for integrating omics data with textual biological knowledge. Yet there is still no accessible, standardized framework that enables systematic comparison of omics representations with different text encoders within a unified workflow. We present mmContext, a lightweight and extensible multimodal embedding framework built on top of the open-source Sentence Transformers library. The software allows researchers to train or apply models that jointly embed omics and text data using any numeric representation stored in an AnnData .obsm layer and any text encoder available in Hugging Face. mmContext supports integration of diverse biological text sources and provides pipelines for training, evaluation, and data preparation. We train and evaluate models for a RNA-Seq and text integration task, and demonstrate their utility through zero-shot classification of cell types and diseases across four independent datasets. By releasing all models, datasets, and tutorials openly, mmContext enables reproducible and accessible multimodal learning for omics–text integration.

**Availability and implementation:** Pretrained checkpoints and full source code for our custom MMContextEncoder are available on Hugging Face huggingface.co/jo-mengr. The Python package github.com/mengerj/mmcontext provides the model implementation and training and evaluation scripts for custom training.

## Introduction

The large body of annotated omics datasets, together with extensive biological knowledge, provides rich context for understanding cellular states. Recent work (Schaefer et al., 2025) on RNA-seq data has shown that such contextual information can be integrated by embedding textual descriptions of cells or samples using language models pre-trained on biomedical literature (Turc et al., 2019; Lee et al., 2020). Aligning these textual embeddings with gene-expression representations produces a joint latent space that combines complementary biological structure encoded in each modality, resulting in a more informative representation.

Existing implementations of such multimodal omics/text models remain difficult to customize and are not integrated into widely supported ecosystems such as Hugging Face. While foundation-model approaches are now used across omics domains, including single-cell and bulk transcriptomics, proteomics, and metabolomics, to learn generalizable biological representations (Chen and Zou, 2024; Cui et al., 2024; Theodoris et al., 2023; Rosen et al., 2024; Hao et al., 2024; Jumper et al., 2021; Hayes et al., 2025), there is currently no framework that enables direct comparison of different numeric and text-based representations of omics data within a single workflow.

To address these limitations, we introduce mmContext, a flexible multimodal embedding framework built on top of the Sentence Transformers library (Reimers and Gurevych, 2019). The framework allows users to train or apply joint text–omics embedding models, using any numeric representation stored in an AnnData.obsm layer (Virshup et al., 2024) and any text encoder available on Hugging Face. Leveraging a mature ecosystem, mmContext inherits compatibility with diverse training regimes, multi-task learning, and downstream tools that rely on Sentence Transformer embeddings. mmContext enables systematic evaluation of different omics encoders, including Geneformer (Theodoris et al., 2023), scVI (Lopez et al., 2018), PCA, or gene-selection–based features. This allows users to tackle open questions in multimodal omics-text integration, such as whether pre-trained models provide more informative initial representations than simpler baselines such as a selection of genes. In addition to numeric encoders, mmContext supports text-based omics encoders that represent omics profiles as ranked feature lists (“cell sentences”), which can be directly embedded with a language model (Rizvi et al., 2025; Levine et al., 2024; Chen and Zou, 2024; Krissmer et al., 2025). To illustrate the capabilities of our framework, we focus on aligning RNA-seq and textual data for zero-shot cell type and disease annotation, a challenging task where most existing methods rely on integrating test data into a reference dataset or applying a pre-trained classifier (Pasquini et al., 2021), and therefore cannot predict labels unseen during training. In mmContext, the language encoder provides a natural mechanism for zero-shot inference by embedding free-text label descriptions in the same latent space as expression profiles. This RNA-seq–based setting facilitates direct comparison with existing multimodal approaches such as CellWhisperer (Schaefer et al., 2025), while the framework remains broadly applicable to other omics–text integration tasks. We benchmark multiple model configurations, trained on large-scale pseudo-bulk and bulk RNA-seq data, systematically comparing pretrained omics embeddings with simpler numeric representations.

By releasing pretrained models and datasets on Hugging Face, mmContext provides a reproducible, open resource for multimodal exploration of single-cell and bulk transcriptomic data.

### Functionality

The mmContext framework provides a modular interface for training and applying multimodal embedding models that align structured omics features with text descriptions in a shared latent space. Implemented as a custom model within the Sentence Transformers framework (Reimers and Gurevych, 2019), it employs two modality-specific towers to encode omics and text data and optimizes similarity between true pairs while minimizing similarity between mismatched ones. Different dataset types and contrastive-learning objectives available in Sentence Transformers can be used without modification. An overview of the training and evaluation workflow is shown in Figure 1.

**Fig. 1:**
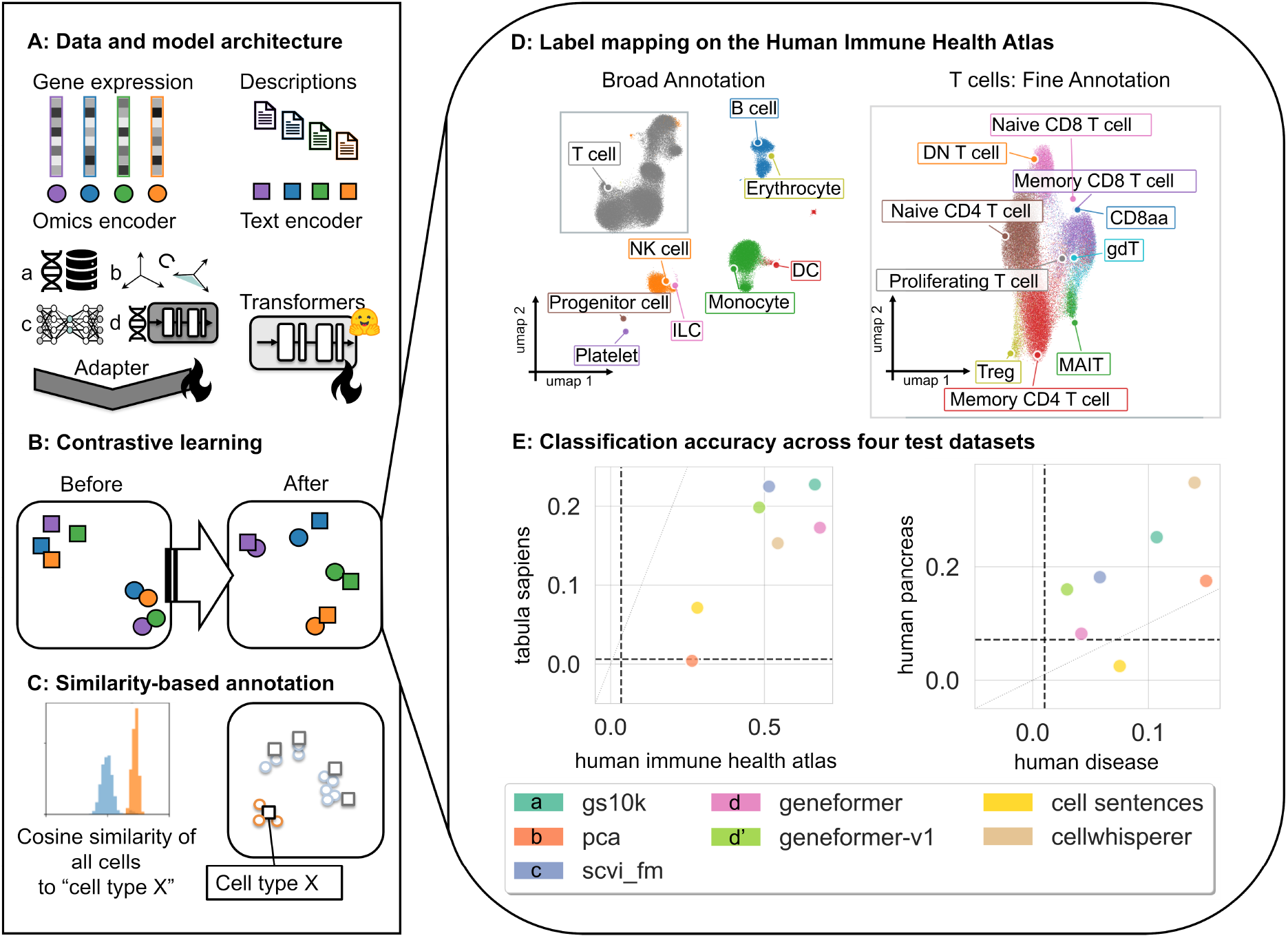
Overview of training and evaluation in mmContext. **(A) Data and model architecture**. Pairs of gene-expression profiles and corresponding text descriptions are passed through two modality-specific encoders. For the omics data, different encoders can be used to generate an initial numeric representation, which is optimized via an adapter layer. Currently implemented are: (a) a gene-selection embedder, (b) principal components of the selected genes, (c) embeddings from a pre-trained scVI model, and (d and d’) embeddings from a pre-trained Geneformer model with 104M parameters and 30M parameters. The text encoder can be any transformer model available on the Hugging Face Hub. (B) **Contrastive learning**. Omics and text modalities are aligned through a contrastive loss that increases similarity between matching pairs and decreases it for non-matching pairs. (C) **Similarity-based annotation**. After training, any text query (e.g., a cell-type label) can be embedded into the shared space to identify the most similar omics samples by cosine similarity. (D) **Label mapping on the Human Immune Health Atlas**. UMAP projection of the joint latent space shows well-aligned broad and fine-grained immune cell annotations on unseen test data. Created with the gs10k model, on a subset of 100k cells. (E) **Classification accuracy across four test datasets**. Each point represents the accuracy on two datasets achieved by a certain model; the dotted line indicates the random baseline (1*/n*_classes_) accuracy per dataset. Besides the four initial omics encoders shown in A, a model using “cell sentences”, where count data is transformed into a list of gene names, is trained and evaluated with the mmContext framework. These mmContext models are compared to the CellWhisperer model by Schaefer et al. (2025).

Although this work demonstrates the approach using RNA-seq data, the same architecture can be directly extended to other omics modalities that can be represented as numeric feature vectors.

Below we outline the training workflow and resources available for working with the mmContext model: (A) Create initial omics representations and descriptions for each sample and store them in an AnnData object. (Virshup et al., 2024) Currently four omics encoders are implemented, which are detailed in the supplementary material. The pairs are used to create a Hugging Face dataset, suitable for training with the Sentence Transformers Trainer. Tutorials and a complete pipeline for these steps are provided at github.com/mengerj/adata hf datasets. The workflow can be run locally or on an SSH server and supports out-of-memory chunking for large datasets. Users can also extend the pipeline by adding their own embedding method via a custom “InitialEmbedder”. Detailed instructions are available in the repository README. (B) Train the mmContext model with a contrastive learning objective, aligning pairs of omics representation and description. (C) Use the trained model to embed a new dataset and query it with free text. Any free-text query can be embedded in the joint space to compute similarity scores against the omics embeddings, enabling flexible analyses such as text-guided exploration or zero-shot classification. For classification, provide a set of candidate labels, embed each label string, compute pairwise similarities, and assign each sample the label with the highest score.

Prior to generating the initial embeddings, all datasets undergo quality control, normalization and log-transformation, following standard single-cell workflows (Luecken and Theis, 2019), including filtering of low-quality cells and genes and batch-aware selection of highly variable genes. These preprocessing steps can be executed automatically before embedding generation and dataset conversion.

Because mmContext inherits the full Sentence Transformers API, users can easily integrate any text encoder available on the Hugging Face Hub (e.g., PubMedBERT, all-MiniLM, or Qwen-Embedding). Additional text-based datasets, such as NCBI abstracts (Krissmer et al., 2025), ontology definitions, or cell-type descriptions, can be incorporated to enrich the semantic structure of the joint embedding space and improve biological context alignment.

To demonstrate the framework’s capabilities, we trained mmContext with large-scale pseudo-bulk and bulk RNA-seq datasets that were also used in the CellWhisperer project, facilitating comparability. We showcase how different initial representations can be combined within the same workflow and evaluated on independent benchmarks for zero-shot cell-type and disease classification (Figure 1 D,E).

### Application Example

We trained mmContext using the pseudo-bulk CellxGene dataset (approx. 20 million cells aggregated into 0.3 million pseudo-bulk profiles) and half the samples from the GEO bulk RNA-seq dataset (approx. 0.35 million samples) curated by Schaefer et al. (2025). Natural-language captions for each sample were generated from metadata, titles, and abstracts using a large language model as described in their work. For evaluation, we used four independent datasets not included in training: (i) the Human Immune Health Atlas with hierarchical cell-type annotations (Gong et al., 2024; Gustafson et al.), (ii) Tabula Sapiens v1 containing over 150 human cell types (Consortium* et al., 2022), (iii) the Human Pancreas single-cell dataset with pronounced batch effects (Luecken and Theis, 2019; Grün et al., 2016; Baron et al., 2016; Lawlor et al., 2017; Xin et al., 2016), and (iv) a bulk RNA disease dataset with more than 100 disease labels (Schaefer et al., 2025).

All test data were unseen during training. For the experiments shown, we used PubMedBERT as the text encoder, producing 768-dimensional embeddings. (NeuML, 2023) Both omics and text encoders were mapped into a shared 2048-dimensional space via two separate two-layer networks. Training employed a contrastive loss (MultipleNegativesRankingLoss) (Henderson et al., 2017) on paired pseudo-bulk profiles and text captions.

The resulting joint embedding space (Figure 1 D), obtained with the gs10k-based model, demonstrates good alignment between textual labels and cell-type embeddings, even for T-cell subtypes. Zero-shot classification was performed by comparing each test sample’s embedding to all label embeddings using cosine similarity, with predictions assigned by nearest-neighbor matching (Figure 1 C). Classification accuracy (Figure 1 E) served as the primary evaluation metric. Despite the difficulty of multi-class datasets such as Tabula Sapiens, the best mmContext model (gs10k) achieved performance over 35-fold above random baseline. Across all benchmarks, models initialized with selected-gene embeddings (gs10k) consistently outperformed those based on pretrained omics embeddings, demonstrating that biologically informed yet lightweight representations can rival large foundation-model encoders. These results illustrate how mmContext enables reproducible comparison of diverse text-omics alignment strategies within a single modular framework. As a complementary measure of ranking quality, the gs10k model achieved an AUC of 0.98 for broad immune lineages and 0.92 for fine grained annotations.

Across the Tabula Sapiens and Human Immune Health Atlas benchmarks, most models achieved comparable performance, with only the PCA-based representation and the cell-sentence approach showing markedly lower accuracy. Interestingly, the PCA model performed best on the human disease dataset, which consists of bulk RNA-seq samples. Combined with its weak performance on single-cell data, this suggests that the PCA embedding captures variation relevant for bulk profiles but does not generalize well across single-cell and bulk domains. The CellWhisperer model showed consistently strong performance across datasets, particularly on the Human Pancreas dataset, which exhibits substantial batch effects.

All datasets and pretrained models used in this evaluation are publicly available on the Hugging Face Hub (https://huggingface.co/jo-mengr), allowing full reproducibility of the presented results and further experimentation within the mmContext framework.

## Conclusion

We introduced mmContext, a practical framework for aligning transcriptomic profiles with natural-language descriptions through contrastive learning built on the Sentence Transformers library. Systematic benchmarking shows that biologically informed gene-selection embeddings provide an effective numeric representation for multimodal alignment, outperforming current pretrained RNA-seq models.

Our findings highlight contrastive multimodal alignment as a viable foundation for zero-shot cell-type classifiers, while revealing remaining challenges in fine-grained subtype recognition. Future developments may include: (i) specialized cell-type models trained exclusively on single-cell data or domain-specific subsets such as immune cells; (ii) incorporation of additional omics modalities to construct richer joint embedding spaces for unpaired data; (iii) applying the model as a classifier or guiding signal for generative diffusion models of transcriptomes; and (iv) integration of mmContext into interactive, text-driven exploration tools for single-cell datasets.

By openly releasing both the framework and pretrained models, together with standardized datasets on the Hugging Face Hub, we provide a reproducible and extensible resource for the community and a foundation for future advances in language-aware multimodal omics analysis.

## Data availability

Both the training data and the test data are publicly accessible through Hugging Face huggingface.co/jo-mengr, in a format that can be directly utilized for training with the SentenceTransformer based mmContext model. The training data was originally collected in Schaefer et al. (2025) and our preprocessed versions are stored on Zenodo.

## Funding

This work is funded by the Deutsche Forschungsgemeinschaft (DFG, German Research Foundation) under Germany’s Excellence Strategy CIBSS – EXC-2189 – Project ID 390939984 (JM, SMK, CK, HB), CRC 1597 Small Data – Project-ID 499552394 (SMK,

CK, HB, MH), CRC 1479 OncoEscape – Project-ID 441891347 (HB), and TRR 359 PILOT – Project-ID 491676693 (HB).

## Author contributions statement

## References

M. Baron, A. Veres, S. L. Wolock, A. L. Faust, R. Gaujoux, A. Vetere, J. H. Ryu, B. K. Wagner, S. S. Shen-Orr, A. M. Klein, et al. A single-cell transcriptomic map of the human and mouse pancreas reveals inter-and intra-cell population structure. Cell systems, 3(4):346–360, 2016.

Y. Chen and J. Zou. Genept: a simple but effective foundation model for genes and cells built from chatgpt. bioRxiv, pages 2023–10, 2024.

T.T.S. Consortium*, R. C. Jones, J. Karkanias, M. A. Krasnow, A. O. Pisco, S. R. Quake, J. Salzman, N. Yosef, B. Bulthaup, P. Brown, et al. The tabula sapiens: A multiple-organ, single-cell transcriptomic atlas of humans. Science, 376(6594):eabl4896, 2022.

H. Cui, C. Wang, H. Maan, K. Pang, F. Luo, N. Duan, and B. Wang. scgpt: toward building a foundation model for single-cell multi-omics using generative ai. Nature Methods, 21(8): 1470–1480, 2024.

Q. Gong, M. Sharma, E. L. Kuan, M. C. Glass, A. Chander, M. Singh, L. T. Graybuck, Z. J. Thomson, C. M. LaFrance, S. R. Zaim, T. Peng, L. Y. Okada, P. C. Genge, K. E. Henderson, E. M. Dornisch, E. D. Layton, P. J. Wittig, A. T. Heubeck, N. M. Mukuka, J. Reading, C. R. Roll, V. Hernandez, V. Parthasarathy, T. J. Stuckey, B. Musgrove, E. Swanson, C. Lord, M. D. A. Weiss, C. G. Phalen, R. R. Mettey, K. J. Lee, J. B. Johanneson, E. K. Kawelo, J. Garber, U. Krishnan, M. Smithmeyer, E. John Wherry, L. Vella, S. E. Henrickson, M. S. Kopp, A. K. Savage, L. A. Becker, P. Meijer, E. M. Coffey, J. J. Goronzy, C. Speake, T. F. Bumol, A. W. Goldrath, T. R. Torgerson, X.-J. Li, P. J. Skene, J. H. Buckner, and C. E. Gustafson. Longitudinal multi-omic immune profiling reveals age-related immune cell dynamics in healthy adults. bioRxiv, page 2024.09.10.612119, Sept. 2024.

D. Grün, M. J. Muraro, J.-C. Boisset, K. Wiebrands, A. Lyubimova, G. Dharmadhikari, M. van den Born, J. Van Es, E. Jansen, H. Clevers, et al. De novo prediction of stem cell identity using single-cell transcriptome data. Cell stem cell, 19 (2):266–277, 2016.

C. E. Gustafson, P. J. Skene, A. W. Goldrath, X.-J. Li, T. R. Torgerson, L. A. Becker, T. F. Bumol, A. Chander, E. M. Coffey, E. M. Dornisch, J. Garber, P. C. Genge, M. Glass, Q. Gong, K. E. Henderson, V. Hernandez, A. T. Heubeck, J. B. Johanneson, E. K. Kawelo, M. S. Kopp, U. Krishnan, E. L. Kuan, C. M. LaFrance, E. D. Layton, K. J. Lee, C. Lord, R. R. Mettey, N. M. Makuka, B. Musgrove, L. Y. Okada, V. Parthasarathy, T. Peng, C. G. Phalen, S. R. Zaim, J. Reading, C. R. Roll, M. Sharma, M. Singh, T. J. Stuckey, E. Swanson, Z. J. Thomson, M. D. A. Weiss, P. J. Wittig, J. H. Buckner, M. Smithmeyer, C. Speake, S. Henrickson, L. Vella, E. J. Wherry, Y. Aggoune, M. Ambrose, A. Beaubien, J. Harvey, N. Howard, N. Inala, E. Johnson, A. Kelsey, M. Kinsey, J. Liang, P. Mariz, S. Pister, S. Subramanian, V. Tereschenko, A. Vetto, P. Meijer, and L. T. Graybuck. AIFI immune health atlas. https://apps.allenimmunology.org/aifi/resources/imm-health-atlas/.

M. Hao, J. Gong, X. Zeng, C. Liu, Y. Guo, X. Cheng, T. Xu, and L. Song. scfoundation: large scale foundation model on single-cell transcriptomics. bioRxiv, 2024.

T. Hayes, R. Rao, H. Akin, N. J. Sofroniew, D. Oktay, Z. Lin, R. Verkuil, V. Q. Tran, J. Deaton, M. Wiggert, et al. Simulating 500 million years of evolution with a language model. Science, 387(6736):850–858, 2025.

M. Henderson, R. Al-Rfou, B. Strope, Y.-H. Sung, L. Lukács, R. Guo, S. Kumar, B. Miklos, and R. Kurzweil. Efficient natural language response suggestion for smart reply. arXiv preprint 1705.00652, 2017.

J. Jumper, R. Evans, A. Pritzel, T. Green, M. Figurnov, O. Ronneberger, K. Tunyasuvunakool, R. Bates, A. Žídek, A. Potapenko, et al. Highly accurate protein structure prediction with alphafold. nature, 596(7873):583–589, 2021.

S. M. Krissmer, J. Menger, J. Rollin, T. M. Vogel, H. Binder, and M. Hackenberg. Adding layers of information to scrna-seq data using pre-trained language models. bioRxiv, pages 2025–08, 2025.

N. Lawlor, J. George, M. Bolisetty, R. Kursawe, L. Sun, V. Sivakamasundari, I. Kycia, P. Robson, and M. L. Stitzel. Single-cell transcriptomes identify human islet cell signatures and reveal cell-type–specific expression changes in type 2 diabetes. Genome research, 27(2):208–222, 2017.

J. Lee, W. Yoon, S. Kim, D. Kim, S. Kim, C. H. So, and J. Kang. Biobert: a pre-trained biomedical language representation model for biomedical text mining. Bioinformatics, 36(4):1234–1240, 2020.

D. Levine, S. A. Rizvi, S. Lévy, N. Pallikkavaliyaveetil, D. Zhang, X. Chen, S. Ghadermarzi, R. Wu, Z. Zheng, I. Vrkic, et al. Cell2sentence: teaching large language models the language of biology. BioRxiv, pages 2023–09, 2024.

R. Lopez, J. Regier, M. B. Cole, M. I. Jordan, and N. Yosef. Deep generative modeling for single-cell transcriptomics. Nature methods, 15(12):1053–1058, 2018.

M. D. Luecken and F. J. Theis. Current best practices in single-cell rna-seq analysis: a tutorial. Molecular systems biology, 15 (6):e8746, 2019.

NeuML. Pubmedbert embeddings. https://huggingface.co/NeuML/pubmedbert-base-embeddings/tree/main, 2023. Accessed: 2025-11-03.

G. Pasquini, J. E. R. Arias, P. Schäfer, and V. Busskamp. Automated methods for cell type annotation on scrna-seq data. Computational and Structural Biotechnology Journal, 19:961–969, 2021.

N. Reimers and I. Gurevych. Sentence-bert: Sentence embeddings using siamese bert-networks. In Proceedings of the 2019 Conference on Empirical Methods in Natural Language Processing. Association for Computational Linguistics, 11 2019. URL http://arxiv.org/abs/1908.10084.

S. A. Rizvi, D. Levine, A. Patel, S. Zhang, E. Wang, S. He, D. Zhang, C. Tang, Z. Lyu, R. Darji, et al. Scaling large language models for next-generation single-cell analysis. bioRxiv, pages 2025–04, 2025.

Y. Rosen, Y. Roohani, A. Agarwal, L. Samotorcan, T. S. Consortium, S. R. Quake, and J. Leskovec. Universal cell embeddings: A foundation model for cell biology. bioRxiv, 2024.

M. Schaefer, P. Peneder, D. Malzl, S. D. Lombardo, M. Peycheva, J. Burton, A. Hakobyan, V. Sharma, T. Krausgruber, C. Sin, et al. Multimodal learning enables chat-based exploration of single-cell data. Nature Biotechnology, pages 1–11, 2025.

C. V. Theodoris, L. Xiao, A. Chopra, M. D. Chaffin, Z. R. Al Sayed, M. C. Hill, H. Mantineo, E. M. Brydon, Z. Zeng, X. S. Liu, et al. Transfer learning enables predictions in network biology. Nature, 618(7965):616–624, 2023.

Turc, M.-W. Chang, K. Lee, and K. Toutanova. Well-read students learn better: On the importance of pre-training compact models. arXiv preprint 1908.08962, 2019.

I. Virshup, S. Rybakov, F. J. Theis, P. Angerer, and F. A. Wolf. anndata: Access and store annotated data matrices. Journal of Open Source Software, 9(101):4371, 2024.

Y. Xin, J. Kim, H. Okamoto, M. Ni, Y. Wei, C. Adler, A. J. Murphy, G. D. Yancopoulos, C. Lin, and J. Gromada. Rna sequencing of single human islet cells reveals type 2 diabetes genes. Cell metabolism, 24(4):608–615, 2016.

