## Supplementary Material for "mmContext: an open framework for multimodal contrastive learning of omics and text data"

### S1 Supplementary Material

#### S1.1 Preprocessing

The training data consists of a collection of publically available (pseudo-)bulk RNA sequencing profiles. This data was gathered and annotated with natural language descriptions by Schaefer et al. (2024). They gathered roughly 350 thousand pseudo-bulk samples from the cellxgene database (CZI Cell Science Program et al., 2025), by taking the mean expression of cells from the same biological context. The second dataset comprises bulk RNA-seq profiles from the NCBI GEO database (Barrett et al., 2005). All natural language descriptions were created with an LLM being prompted with metadata, available titles and abstracts (see Schaefer et al. (2024) for details). The raw h5ad-files are split into chunks to allow processing with limited memory usage. Quality control metrics are computed on each chunk, and low-quality cells are removed. Quality control is performed according to (Luecken et al., 2019), where a mean absolute deviation (MAD) is computed for the following metrics: total counts (log transformed), number of genes (log transformed), and the percent of counts in the top 20 % of genes. Cells with more than 3 MADs difference to the median percent of mitochondrial counts or more than 8% of mitochondrial genes are also removed. Genes present in less than 20 cells are removed and cells with less then 200 non-zero genes are also removed. Cells from low-frequency categories of interest (e.g., batch or cell type) with fewer than five occurrences in the dataset were excluded. To improve the models ability to correct strong batch effects, originating from different sequencing platforms, the instruments used were added to the caption of each cell. Raw counts are log-transformed and normalized. Batch aware highly variable gene selection is performed. Batches are considered to be different studies. Since a strong bimodal split was observed for the amount of aligned reads within the RNA-seq dataset gathered from the GEO database, and this effect dominated the variance of the dataset, we decided to split the dataset according to this bimodal distribution. Each split was then normalized separately. The origin of the split could not be resolved with the available metadata, but it is likely a technical artifact and not a biological one.

#### S1.2 Implementation Details

The Sentence Transformers framework (Reimers et al., 2019) trains dual-encoder models to learn similarity-based embeddings using paired data, maximizing cosine similarity for positive pairs and minimizing it for negatives. For textual input, a tokenizer maps text to token IDs, and a lookup table converts these IDs to embedding vectors.

To enable multimodal learning with numerical omics data, mmContext extends this architecture by introducing a modality-specific mechanism for registering continuous feature vectors (such as gene, protein, or metabolite profiles) as "tokens." Users provide a table that maps unique string identifiers to initial omics embeddings, from which the model constructs a frozen embedding matrix analogous to a text token embedding table. Each identifier can correspond to a cell, sample, or other omics entity, and the associated numeric representation may be defined at either the sample level (e.g., selected genes, PCA features, or embeddings from pretrained RNA-seq models) or the feature level (e.g., a protein-language-model embedding of the amino-acid sequence of a transcript, gene, or protein).

Because these omics embeddings are continuous and sample-specific rather than drawn from a fixed vocabulary, the model cannot rely on a persistent token dictionary as is done for text. Instead, the corresponding embeddings must be supplied whenever a new dataset is used. In practice, initial embeddings are stored in an AnnData object (Virshup et al., 2024) and loaded into the model at inference time through the mmContext API, ensuring that the embedding table is consistent with the method used during training.

At runtime, all inputs are handled as strings annotated with modality-specific prefixes. When a string begins with a registered omics prefix, the omics tokenizer retrieves the correct initial embedding from the frozen embedding matrix. Text strings are processed with the chosen transformer tokenizer as usual. This design preserves full compatibility with the Sentence Transformers input pipeline while enabling fast lookup of numeric omics embeddings and seamless integration of mixed omics-text inputs within a shared latent space.

The currently implemented initial embedders are described in the following. (a) Gene selection (gs10k). Uses the expression matrix of 10,000 selected genes, including all known marker genes from CellMarker and PanglaoDB (details below). Genes absent from a dataset are set to zero. (b) Principal components. Obtained by applying PCA to the selected genes, reducing dimensionality to 50. The transformation is fitted jointly on subsets of the training data from CellxGene and GEO (roughly 80k samples from each dataset)(CZI Cell Science Program et al., 2025; Barrett et al., 2005), and the same loadings are reused for test data to ensure consistency between training and evaluation. (c) scVI embeddings. Derived from a pre-trained scVI model trained on the CellxGene corpus, producing a 50-dimensional embedding space (Ergen et al., 2025). In the following, this representation is referred to as scvi_fm (foundation model). (d) Geneformer embeddings. Obtained from the transformer-based Geneformer model (Theodoris et al., 2023), pre-trained on approximately 104 million cells and comprising 104 million parameters. The generated cell embeddings have 768 dimensions. Additionally, the Geneformer-v1 was used (d'), derived from the first version of the Geneformer model used in the CellWhisperer project, trained on approximately 30 million cells and producing 512-dimensional embeddings.

As an additional text-based input representation, cell sentences can be used, obtained by converting each expression profile into a ranked list of the most highly expressed genes which can be directly processed by a text encoder (Rizvi et al., 2025; Levine et al., 2024), jointly with the accompanying textual descriptions.

##### S1.3 Gene Selection

To define a compact but biologically relevant gene set, we combined reference gene annotations from GENCODE v49 (GRCh38.p14) with transcriptomic measurements from the Human Protein Atlas (HPA) consensus RNA dataset. The latter integrates RNA-seq data from HPA and GTEx across 50 consensus tissues, reporting normalized transcript counts (nTPM) for each gene. GENCODE annotations were parsed at the gene level to obtain Ensembl identifiers, HGNC symbols, and biotypes. To ensure compatibility with HPA data, Ensembl gene identifiers were normalized by removing version suffixes (e.g. ENSG00000141510.18 $\rightarrow$ ENSG00000141510). We restricted the gene universe to selected biotypes that are most relevant for transcriptomic analyses: protein coding, lincRNA, antisense, and processed transcript. Genes annotated as ribosomal RNAs (e.g. rRNA, Mt_rRNA), tRNAs, or pseudogenes were excluded. From the HPA consensus file (rna_tissue_consensus.tsv.zip), we generated a gene × tissue expression matrix using nTPM values. For genes with multiple entries per tissue, the maximum nTPM was retained, consistent with the consensus definition. Genes were required to be expressed above a detection threshold ($\geq$1 nTPM) in at least five tissues to minimize inclusion of spurious low-abundance transcripts. For each gene, three statistics were computed across the 50 tissues: 1. Mean expression ($\mu$) 2. Variance of expression ($\sigma^2$) 3. Tissue specificity (Tau), defined as

$$\tau = \frac{\sum_{i=1}^{n} (1 - x_i / \max_j x_j)}{n - 1} \quad (1)$$

where $x_i$ denotes expression in tissue i. Tau ranges from 0 (ubiquitous expression) to 1 (restricted to a single tissue).

Each statistic was standardized (z-scored) across all genes, and a composite relevance score was calculated as

$$\text{score} = 0.5 \cdot z(\sigma^2) + 0.4 \cdot z(\tau) + 0.1 \cdot z(\mu). \quad (2)$$

This weighting prioritizes genes with high variability and tissue-specific patterns, while retaining moderately expressed ubiquitous genes.

Genes were ranked by the composite score and the top 10,000 were retained as the analysis panel. To ensure inclusion of well-established marker genes (e.g. immune and developmental markers), we additionally unioned curated marker sets from CellMarker and PanglaoDB with the ranked list. Duplicate entries were resolved by gene symbol, and the resulting panel contained both protein-coding genes and non-coding RNAs with established or putative biological roles.

This approach balances breadth and relevance: it avoids the inflation caused by tens of thousands of non-coding loci in GENCODE, yet preserves tissue-specific and regulatory genes (including lncRNAs) that may be critical for cell identity and disease.

#### S1.4 Training Details

After processing the training data, including the creation of the initial embeddings, datasets were stored on huggingface. These datasets contain both a cell token, which is simply the cell identifier, a cell sentences of 4096 genes, which can be truncated at training time and a path poiniting to the adata object, which can also be a share link to a cloud store. For all shown models, training is performed for 16 epochs with a batch size of 512, while keeping the text encoder frozen for the first epoch, thus only optimizing the adapter layers. The MultipleNegativesRankingLoss from the SentenceTransformers library is used, including two hard negatives, representing different cell types from the same batch. (Henderson et al., 2017) All shown models were trained with the PubMedBert-Embeddings model, which we found to work well for this task, while being computationally light. (NeuML, 2023) The training parameters, including the output dimension, were mainly chosen to match those used to train the CellWhisperer model, in order to facilitate comparison.

#### S1.5 Usage of LLMs

Large Language models were utilized to write and improve code and documentation as well as to improve the manuscript.
